# Neurocomputational mechanisms underlying motivated seeing

**DOI:** 10.1101/364836

**Authors:** Yuan Chang Leong, Brent L. Hughes, Yiyu Wang, Jamil Zaki

**Affiliations:** Department of Psychology, Stanford University; Department of Psychology, University of California, Riverside; Department of Psychology, Northeastern University

**Keywords:** Motivation, perceptual decision-making, computational modeling, striatum, ventral temporal cortex

## Abstract

People tend to believe their perceptions are veridical representations of the world, but also commonly report perceiving what they want to see or hear, a phenomenon known as *motivated perception*. It remains unclear whether this phenomenon reflects an actual change in what people perceive or merely a bias in their responding. We manipulated the percept participants wanted to see as they performed a visual categorization task for reward. Even though the reward maximizing strategy was to perform the task accurately, this manipulation biased participants’ perceptual judgments. Motivation increased activity in voxels within visual cortex selective for the motivationally relevant category, indicating a bias in participants’ neural representation of the presented image. Using a drift diffusion model, we decomposed motivated seeing into response and perceptual components. Response bias was associated with anticipatory activity in the nucleus accumbens, whereas perceptual bias tracked category-selective neural activity. Our results highlight the role of the reward circuitry in biasing perceptual processes and provide a computational description of how the drive for reward can lead to inaccurate representations of the world.

---

People tend to think of their perception as a veridical representation of the external world, but this view has long been challenged by psychological research^1,2^. Instead, people often report percepts that they are motivated to perceive, a phenomenon we term *motivated perception*. In one classic example in the visual domain, Dartmouth and Princeton students watched the same football game. Fans of each team subsequently reported seeing the other team commit more fouls^3^. Likewise, participants presented with ambiguous line drawings were more likely to report seeing the interpretation associated with desirable outcomes^4^.

One interpretation of these findings is that motivational factors, such as desires and wants, exert top-down influence over perceptual processing, such that people become biased towards seeing what they want to see^5^. We refer to the bias in perceptual processing as a *perceptual bias*. Alternatively, these effects could instead reflect a *response bias:* a bias not in what participants see, but merely in what they report seeing^6,7^. Although these two interpretations appear at odds with each other, they are not mutually exclusive; motivation could simultaneously both bias perception and responses. Computational models offer a promising analytical approach by which we can dissociate these two sources of bias and identify their independent contributions to perceptual judgments.

Drift diffusion models assume that perceptual judgments arise from the accumulation of noisy sensory evidence towards one of two decision thresholds^8,9^. When the level of evidence exceeds the threshold associated with a particular percept, the corresponding response is made. Within this framework, a response bias can be modeled as a bias in the *starting point* of evidence accumulation. This reduces the amount of evidence needed to make a response, but assumes no effect on perceptual processing. On the other hand, a perceptual bias can be modeled as a bias in the *rate* of evidence accumulation. This in turn reflects sensory information accumulating faster for one percept than another, implying that perceptual processes are biased towards seeing that percept. The extent to which each bias influences behavior can then be estimated from empirical data.

Neuroimaging offers a second, complementary approach through which to dissociate response and perceptual biases. The neural mechanisms underlying motivational effects on perceptual judgments are not well understood, but separate literatures on the neuroscience of motivation and perception suggest distinct neural processes that could be related to different components of bias. In particular, both fMRI and electrophysiology studies have identified the nucleus accumbens (NAcc) as a key structure in mediating motivational processes^10,11^. One putative role of the NAcc is that it biases response selection in favor of actions associated with higher reward^12–14^. Notably, greater NAcc activity precedes approach behavior to reward-predicting stimuli^15^, while inactivating NAcc reduces the preference for responses associated with larger rewards^16^. We thus predict that the NAcc would play a role in response biases by increasing the readiness to make motivationally desirable judgments.

On the other hand, previous work suggests that perceptual judgments are determined by comparing the activity of neurons selective to different perceptual features^17,18^. For example, monkeys in a direction-of-motion task were more likely to categorize a cloud of dots as moving upward when activity was higher in sensory neurons preferring upward motion than in sensory neurons preferring downward motion^19^. Similarly, Heekeren and colleagues demonstrated in humans that perceptual judgments on a face-scene categorization task were computed by comparing activity in areas in the ventral temporal cortex selective to each category^20^. Motivation could potentially bias this comparison process by driving attention towards the features associated with a motivationally desirable percept^21^. This enhances the neural response to those features, thus giving rise to a perceptual bias.

The goal of the present study was two-fold: (i) to decompose motivational influences on perceptual judgments into a response bias and a perceptual bias and (ii) to examine the neurocomputational mechanisms underlying motivational biases on perceptual judgments. Human participants were presented with visually ambiguous images created by morphing a face image and a scene image together, and were rewarded for correctly categorizing whether the face or scene was of higher intensity. We manipulated participants’ motivation by instructing them on each trial that they would win or lose extra money if the upcoming stimulus was of a particular category. Crucially, participants would gain or lose this additional money based only on the actual category of the stimulus, not what they reported seeing. As such, even though participants were motivated to see one category over the other, they would earn the most money on the task if they reported the stimulus category accurately.

We estimated the magnitude of response and perceptual biases exhibited by our participants by fitting a drift diffusion model to choice and reaction time data. Using fMRI, we searched for distinct neural processes associated with each bias. Furthermore, as the perception of faces and scenes is associated with distinct patterns of activity in the ventral occipito-temporal cortex^20–22,23^, we used multivoxel pattern analysis to measure the level of face- and scene-selective activity as a correlate of perception. If the motivation to see one category increases the level of neural activity selective for that category, it would provide additional evidence that motivation modulates perceptual processing. By combining the neural measures with computational modeling, our approach provides a mechanistic account of motivational influences on perceptual judgments.

## Results

Thirty participants were scanned using fMRI while they performed a categorization task with visually ambiguous images comprising a mixture of a face and a scene (Fig. 1A). For each image, participants were rewarded for correctly indicating which category was of higher intensity (i.e. “more face” or “more scene”). To motivate participants to see one category over another, we informed them that they would be performing the task with a teammate or an opponent. This other “player” would bet on whether the upcoming image would be one with more face or more scene. Participants were told that neither the teammate nor opponent had seen the upcoming image and their bets provided no informational value. Participants won a monetary bonus if the teammate’s bet was correct, and lost money if the teammate’s bet was wrong *(Cooperation Condition)*. In contrast, participants lost money if the opponent’s bet was correct, and won a bonus if the opponent’s bet was wrong *(Competition Condition)*. The *Competition Condition* allowed us to assess the effect of motivation above and beyond that of semantic priming due to having seen the words “Face” and “Scene”. Crucially, the outcome of the teammate’s and opponent’s bets were determined by the objective face-scene proportion of the presented image, and not by participants’ subjective categorizations. To earn the most money, participants should ignore the bets and make their categorizations accurately (Fig. 1B).

**Figure 1.**
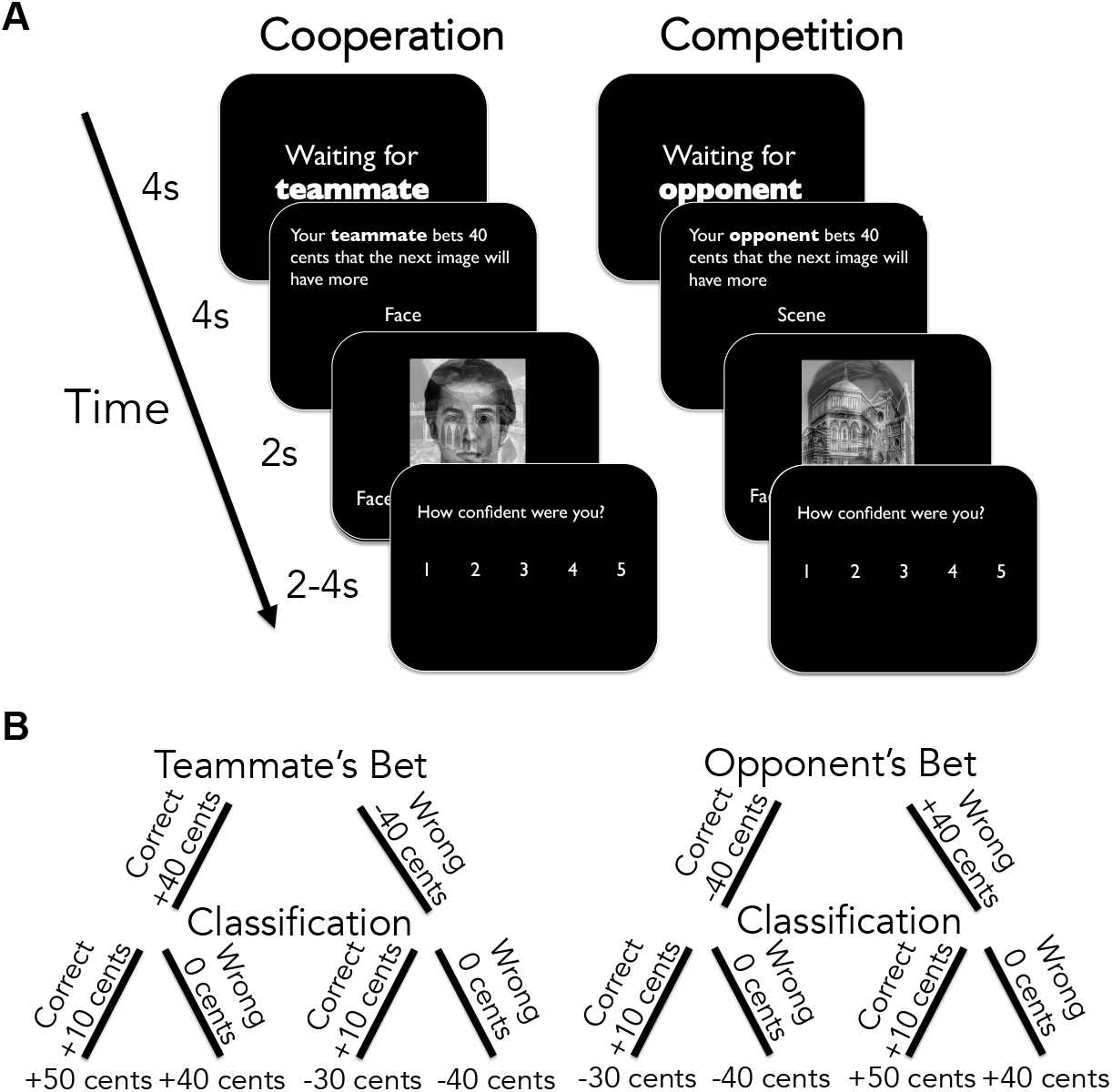
*Experimental Design*. **A.** Motivated Visual Categorization Task. Participants were presented with composite face/scene images. In the Cooperation Condition, a teammate first makes a bet on whether the face or scene will be of higher intensity (i.e. “more face” or “more scene”). Participants are then presented with the composite image, and have to categorize whether it comprises mostly face or mostly scene. They then rated how confident they were in their categorization. In the Competition Condition, an opponent makes the bet instead. Participants performed 2 blocks of each condition, each with 40 trials. All bets were pseudo-randomly generated such that both the teammate and the opponent made correct bets on 50% of the trials. The order of the blocks was interleaved and counterbalanced across participants. **B.** Payoff structure. Participants won an extra 40 cents if the teammate’s bet was correct, but lost 40 cents if the teammate’s bet was wrong. Conversely, they lost 40 cents if the opponent’s bet was correct, but won 40 cents if the opponent’s bet was wrong. Participants earned 10 cents for each correct categorization. As the outcome of the bets was determined by the objective face-scene ratio of the presented image and not by participants’ subjective categorizations, the reward maximizing strategy was to ignore the bets and perform the categorizations accurately.

### Motivation biases visual categorization

For each condition, we estimated the psychometric function describing the relationship between participants’ categorizations and the relative proportions of face and scene in an image. Not surprisingly, as the proportion of scene in an image increases, participants were more likely to categorize the image as having more scene (b = 2.19, SE = 0.13, z = 16.9, p < 0.001; statistical significance was assessed using a generalized linear mixed effects model, see Methods).

To examine the effect of motivation, we estimated separate psychometric functions depending on the teammate or opponent’s bet (Fig. 2A). In the *Cooperation Condition*, participants were more likely to report seeing more scene when the teammate bet on scene than when the teammate bet on face (b = 0.33, SE = 0.13, z = 2.52, p = 0.012). That is to say, participants were more likely to report seeing the category that the bet motivated them to see.

**Figure 2.**
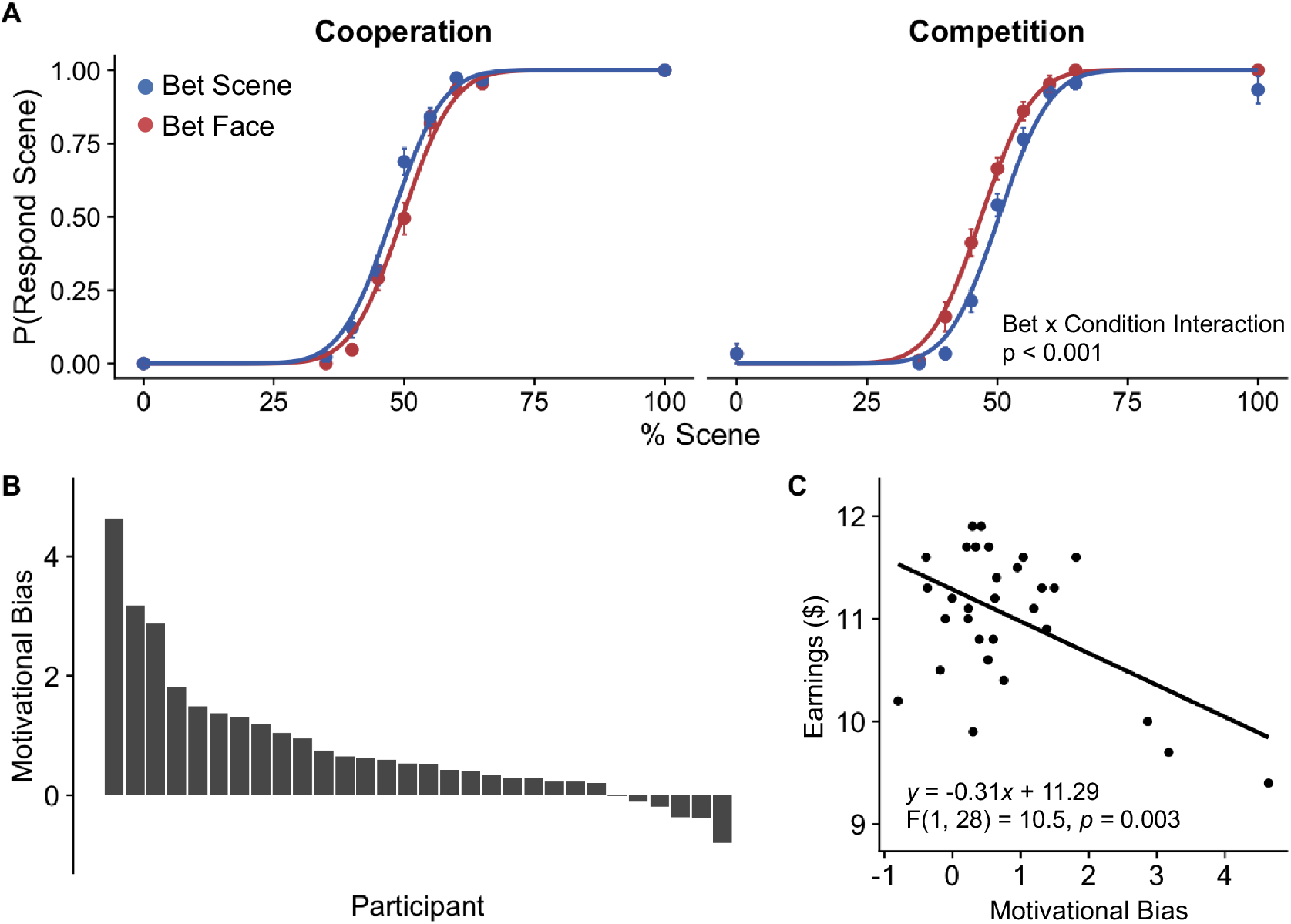
Motivation biases visual categorization. **A.** Participants were more likely to categorize the ambiguous image as what they wanted to see. *Cooperation Condition:* Participants’ psychometric function was shifted left when the teammate bet on more scene (blue) relative to when the teammate bet on more face (red), indicating that less scene evidence is needed to categorize an image as having more scene. *Competition Condition:* Participants’ psychometric function was shifted right when the opponent bet on more scene (blue) relative to when the opponent bet on more face (red), indicating that more scene evidence is needed to categorize an image as having more scene. Statistical significance was assessed using a generalized linear mixed-effects model (see Methods). Error bars indicate S.E.M. **B.** Magnitude of bias in each participant, defined as the random slope of the Bet x Condition interaction. Higher values indicate stronger bias. **C.** Participants with greater motivational bias performed worse on the task and received lower earnings. Statistical significance was assessed using a robust regression analysis that down-weights the influences of outliers. See Fig. S2 for replication with an independent group of participants.

The bias in participants’ perceptual judgments could also be due to semantic priming. For example, when the teammate bet that the upcoming image would have more face, participants might be more likely to report seeing more face because they were semantically primed by having just seen the word “face”, and not because they were motivated to see more face. The *Competition Condition* allows us to directly test this competing account.

In the *Competition Condition*, participants were motivated to see the category that was inconsistent with the opponent’s bet. For example, if the opponent bet that the upcoming image would have more scene, participants would be motivated to see more face. If the bias in participants’ judgments resulted from semantic priming, participants would instead be more likely to report seeing the category consistent with the opponent’s bet. Consistent with a motivational account, participants were *less* likely to categorize an image as having more scene when the opponent bet scene than when the opponent bet face (b = −0.47, SE = 0.11, *z* = −4.11, *p* = 0.012). These results also highlight the flexible nature of the motivational bias, as participants were able to remap the relationship between the word presented to them and the percept they were motivated to see based on the experimental context.

To quantify the magnitude of motivational bias across the two conditions, we computed the *Condition* x *Bet* interaction on participants’ categorizations. This interaction was highly significant (b = 0.81, SE = 0.24, *z* = 3.35, *p* < 0.001), such that participants were more likely to make categorizations consistent with the teammate’s bet and inconsistent with the opponent’s bet. Taken together, these results indicate that participants’ categorizations were biased by what they were motivated to see.

We estimated each participant’s motivational bias by extracting the random slopes of the Condition x Bet interaction. Although the majority of participants exhibited motivational bias, the degree of bias varied across individuals (Fig. 2B). Participants who exhibited stronger motivational bias made fewer correct categorizations, indicating that the motivational bias impaired performance on the task and led to decreased earnings (robust regression: b = −0.31, SE = 0.09, F(1, 28) = 10.5, *p* = 0.003, Fig. 2C). Additional analyses suggest that the performance impairment was not due to general differences in attention or perceptual sensitivity, but because some participants were more affected by the motivation manipulation than others (*Supplemental Note, Fig. S1*). All behavioral findings replicated in a separate group of 28 participants who performed the task without undergoing fMRI (Fig. S2).

### Motivation biases both starting point and drift rate

Having established that participants’ categorizations were biased by what they wanted to see, we proceeded to examine *how* motivation biased the decision process. To this end, we fit a drift diffusion model (DDM) to participants’ choice and reaction time (RT) data. The DDM is a model of the cognitive processes involved in two-choice decisions^9^, and assumes that choice results from the accumulation of noisy sensory evidence towards one of two decision thresholds. The starting point of the accumulation process is determined by a free parameter, z, and the decision threshold is determined by a free parameter, *a*. The rate of evidence accumulation is determined by the drift rate, v, which depends on the sensory information on each trial. In the case of our task, an image with a high scene proportion would be associated with a highly positive *v*, while an image with a high face proportion would be associated with a highly negative *v*. When the accumulation process reaches one of the two thresholds (top threshold for scene, bottom threshold for face), a response is initiated.

From a DDM perspective, our participants’ motivational bias could reflect either or both of two mechanisms (Fig. 3A). First, a shift in the *starting point, z*, could result in an *a priori* bias to make motivation consistent judgments. In particular, shifting the starting point towards the decision threshold of the motivation consistent category reduces the amount of evidence needed to make the motivationally consistent response, thus creating a *response bias*. Second, a bias in the *drift rate, v*, could favor evidence accumulation in favor of the motivationally consistent category. This results in sensory evidence accumulating faster for the motivationally consistent category, thus creating a *perceptual bias*. Both biases increase the proportion of motivation consistent judgments, but have distinguishable effects on the shape of reaction time distributions^9,24^ (Fig. S3).

**Figure 3.**
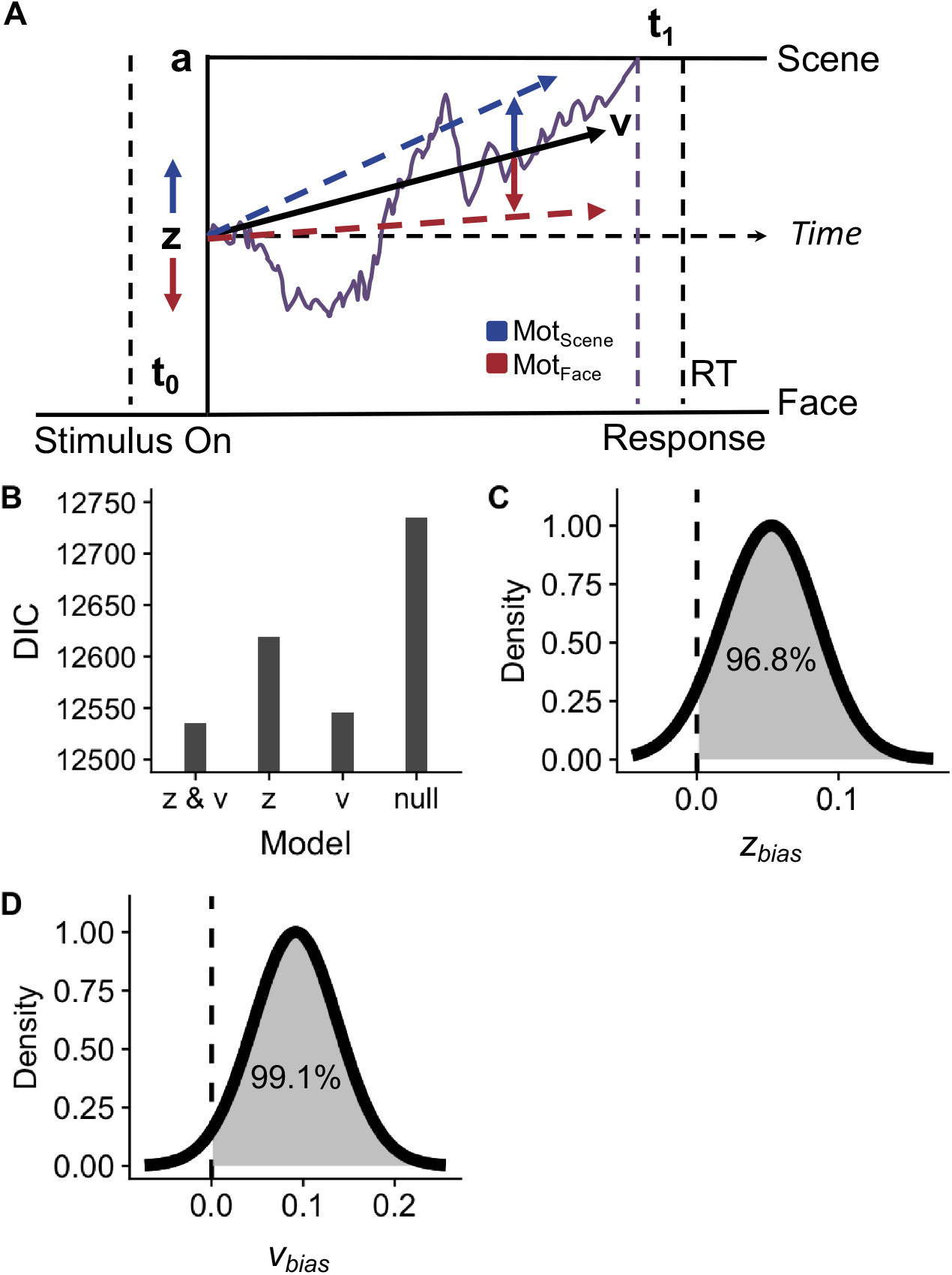
Modeling Results. **A.** Schematic diagram of the drift diffusion model, *t_0_* and *t_1_*. non-decision time related to stimulus encoding and response execution; *a*: decision threshold, *z*: starting point, *v*: drift rate. On each trial, choice depends on the accumulation of noisy sensory evidence towards one of two decision thresholds. Motivation biases categorizations by modulating both the starting point and the drift rate. Blue: Motivated to see more scene; Red: Motivated to see more face. **B.** The *z & v* model has the lowest DIC score, indicating that it provides the best fit to participants’ data. See Table S1 for best-fit values and summary statistics of all model parameters. **C.** Posterior distribution of the starting point bias (*z_biaz_*) Dashed line indicates 0 (no bias). More than 95% of the distribution was greater than 0, indicating credible evidence of a starting point bias. **D.** Posterior distribution of the drift bias (*v_biaz_*). More than 95% of the distribution was greater than 0, indicating credible evidence of a drift bias.

To examine if either or both of these processes explained the bias observed in our task, we fit three different DDMs to participants’ data^25^ (see Methods): i. a model in which motivation biases the starting point (*z*-model), ii. one in which motivation biases the drift rate (*v*-model), iii. and one in which motivation biases both the starting point and drift rate (*z* & *v* model). For comparison, we also fit an unbiased model in which neither the starting point nor drift rate was biased by motivation (*null*-model). As the pattern of reaction times was comparable across the Cooperation and Competition conditions (Fig. S4), all models were fit to data pooled over both conditions to take advantage of the larger number of trials for more reliable estimates.

We compared the model fits based on the deviance information criterion (DIC^26^, a common metric of model comparison for hierarchical models that penalizes for model complexity, with lower values indicating better fit). To verify that DIC is an accurate metric for model comparison, we fit the models to simulated data and demonstrated that DIC reliably identifies the true model used to generate each dataset (*Supplemental Note*, Fig. S5). When the models were fit to experimental data, DIC identified the *z* & *v* model as the model that provided the best fit to participants’data (DIC: *z* & *v*: 12535; *z*: 12619; *v:* 12546, *null:* 12735; Fig. 3B), suggesting that motivation biased both the starting point and drift rate of evidence accumulation. The best-fit values and summary statistics of all model parameters are reported in Table S1.

Next, we examined how the starting point and drift rate were affected by motivation. We extracted the posterior distribution of the starting point bias estimated by the *z & v* model (Fig. 3C). Positive values indicate a positive motivational bias, such that the starting point is biased towards the scene threshold when participants were motivated to see more scene, and biased towards the face threshold when participants were motivated to see more face. The posterior distribution of the starting point bias was credibly positive (*p*(*z_bias_* > 0 = 0.968, M = 0.05, 95% CI [0.00, 0.11]), indicating that motivation biased the starting point of evidence accumulation towards the threshold of the motivation consistent category.

Similarly, the posterior distribution of the bias in drift rate estimated by the *z* & *v* model was credibly positive (*p*(*v_bias_* > 0 = 0.991, M = 0.09, 95% CI [0.02, 0.17], Fig. 3D), indicating that evidence accumulation was biased towards the face category when participants were motivated to see more face, and towards the scene category when participants were motivated to see more scene. Taken together, the modeling results suggest that motivation biased perceptual judgments by increasing the predisposition to respond in a motivation consistent manner, as well as by biasing sensory processing in favor of the motivation consistent category. In later sections, we provide converging neural evidence of this account.

### Drift diffusion model accounts for asymmetries in reaction time

Participants were faster at making categorizations that were consistent with their motivations (*b* = −0.05, *SE* = 0.02, *t*(27) = −2.80, *p* = 0.009). In particular, participants were faster to categorize an image as a face when motivated to see more face (*b* = −0.07, *SE* = 0.02, *t*(28) = −3.12, *p* = 0.004), and faster to categorize an image as a scene when motivated to see more scene (*b* = −0.04, *SE* = 0.02, *t*(28) = −2.11, *p* = 0.04; Fig. 4A). We examined if the *z & v* model would account for this feature of the data. We simulated choice and reaction time data using the *z & v* model (see Methods), and showed that the simulated data reproduced the pattern of reaction times where motivation consistent responses were faster than motivation inconsistent responses (Fig. 4B). We show as well that the model simulations reproduced each participant’s choice and reaction time distributions (Fig. S6). Altogether, these results indicate that model fits of the *z & v* model align well with participants’ data.

**Figure 4.**
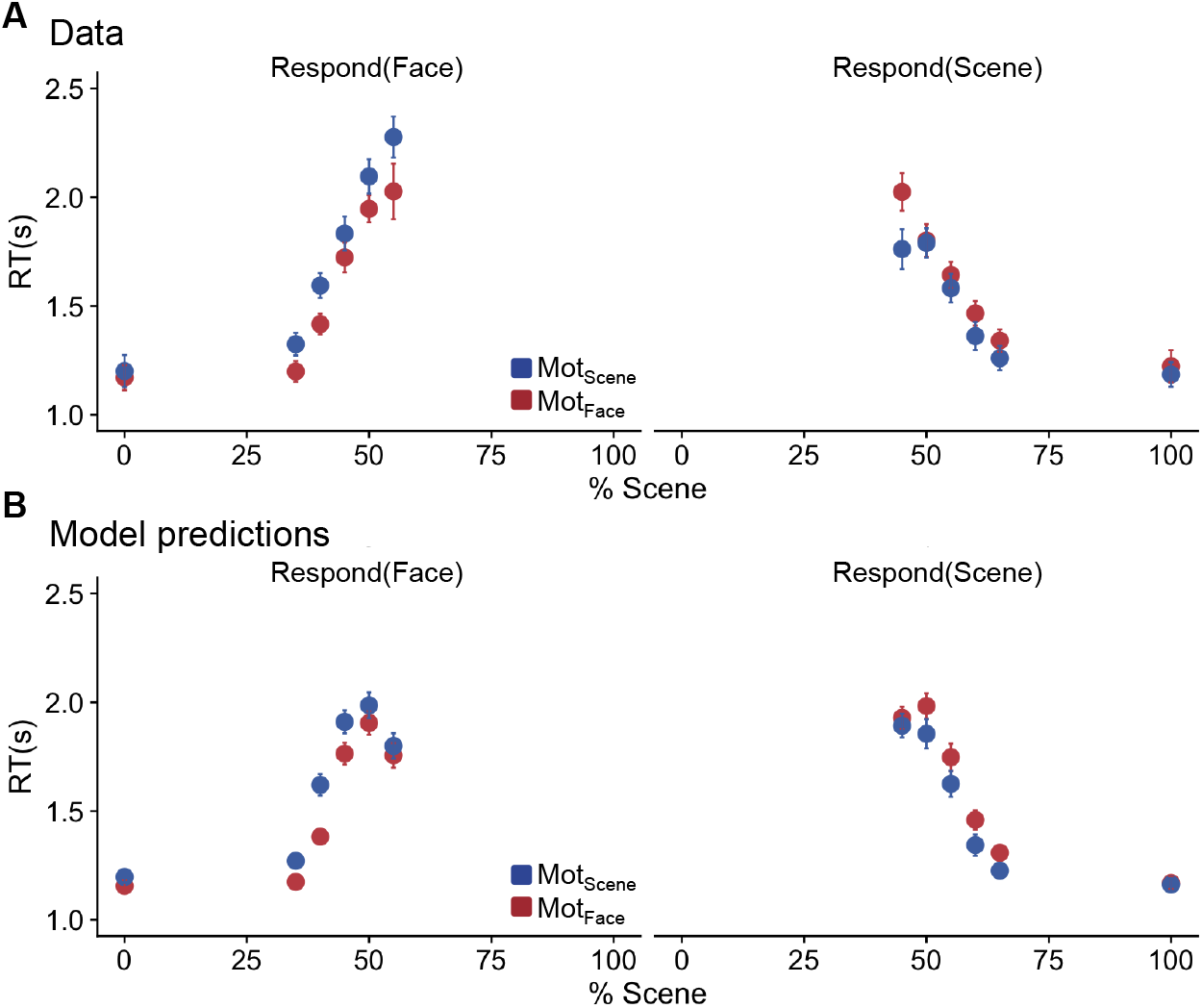
Drift diffusion model accounts for asymmetries in reaction times. **A.** Reaction times for face (left) and scene (right) responses at each % scene, separately for when participants were motivated to see faces (red) and scenes (blue). Participants were faster to categorize an image as the category they were motivated to see. Trial types with less than 48 trials (i.e. 1% of total number of trials) were excluded from the plot because there were too few trials for reliable estimates and they tend to come from a small number of participants. For example, when % scene was high, very rarely did participants categorize an image as a face. Error bars indicate between-subject SEM. **B. Model-predicted reaction times.** Reaction times were simulated using the *z & v* model with parameter values sampled from the posterior distribution. 500 datasets were simulated, each with the same number of participants and trials as the original data. Reaction times were first averaged over simulations to obtain an average reaction time for each trial type for each participant, and then averaged over participants to obtain the mean reaction time for each type. Error bars denote between-subject SEM.

### Motivation consistent categorizations are associated with activity in the salience network and dorsal attention network

To identify the brain areas associated with motivational biases in perceptual judgments, we first performed a whole-brain contrast to identify voxels that responded differently on trials on which participants categorized an image as the category they were motivated to see (*Motivation Consistent* trials) than on trials on which they categorized an image as the category they were motivated to not see (*Motivation Inconsistent* trials). This contrast revealed activations in two network of brain regions: i. the salience network, which includes the nucleus accumbens (NAcc), insula, dorsal anterior cingulate (dACC), and ii. the dorsal attention network, including the intraparietal sulcus (IPS) and frontal eye-fields (FEF) (Fig. 5, https://neurovault.org/collections/EAAXGDRJ/images/62743/)

**Figure 5.**
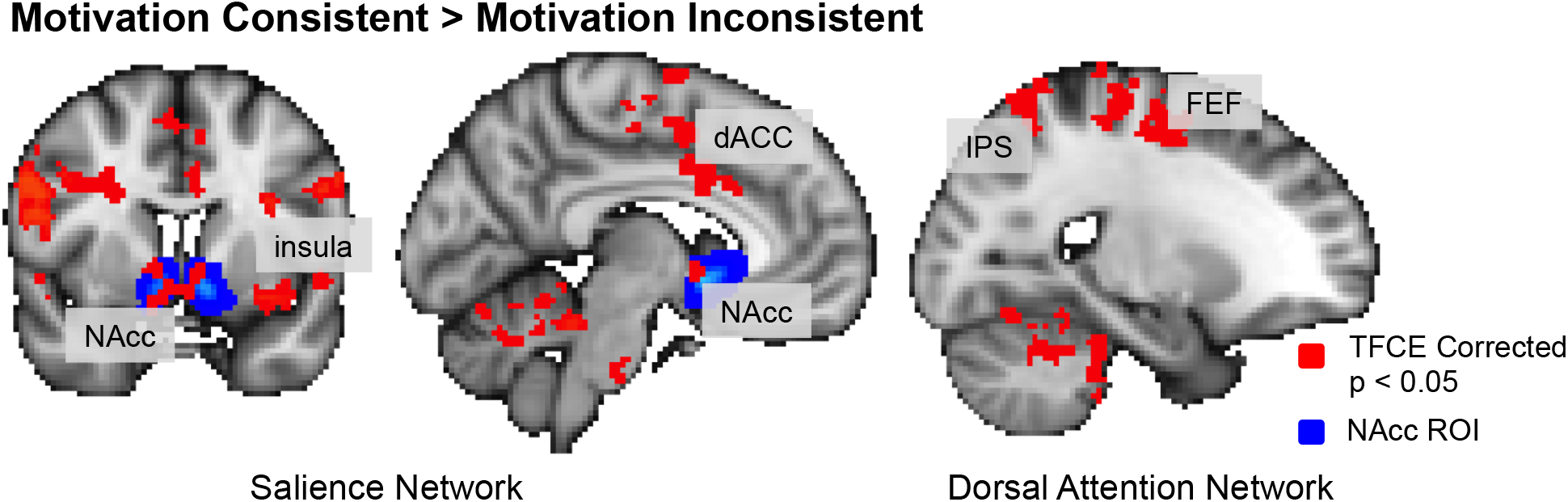
Neural correlates of motivational bias. Activity in the salience and dorsal attention networks was higher when participants made motivation consistent categorizations than when participants made motivation inconsistent categorizations. NAcc: nucleus accumbens, dACC: dorsal anterior cingulate cortex, IPS: intraparietal sulcus, FEF: Frontal eye fields. Correction for multiple comparisons was performed using threshold-free-cluster-enhancement (TFCE) with an alpha of 0.05^27^. Shown in blue is a NAcc ROI defined using the Harvard-Oxford Cortical Structural Atlas.

### NAcc activation is associated with response bias

The NAcc is thought to be crucial in mediating the effects of motivation on actions, and has been previously implicated in biasing responses towards actions associated with larger rewards^12–14^. We thus predicted that the NAcc would be associated with the response bias when making motivated perceptual judgments. We defined a NAcc region of interest (ROI) using the Harvard-Oxford Cortical Structural Atlas, and computed the NAcc response of each participant as the average z-statistic of the Motivation Consistent > Motivation Inconsistent contrast in the ROI (Fig. 5). This value reflects the extent to which the NAcc of a particular participant was more active when they made motivationally consistent categorizations than when they made motivation inconsistent categorizations. We then examined if the NAcc response would be associated with either participants’ response bias, their perceptual bias, or both biases.

For each participant, we computed response bias as the posterior mean estimate of that participant’s starting point bias (*z_bias_*, Fig. 3C), and perceptual bias as the posterior mean estimate of their drift bias (*v_bias_*, Fig. 3D). Estimates of the two biases were not significantly correlated (r = 0.290, p = 0.120). When both biases were entered as predictors in the same linear regression model, participants’ response bias was associated with the NAcc response (*β_z_* = 0.47, SE = 0.17, *t(27)* = 2.71, *p* = 0.012), but their perceptual bias was not (*β_v_*, = 0.07, SE = 0.17, *t(27)* = 0.37, *p* = 0.711, Fig. 6A).

**Figure 6.**
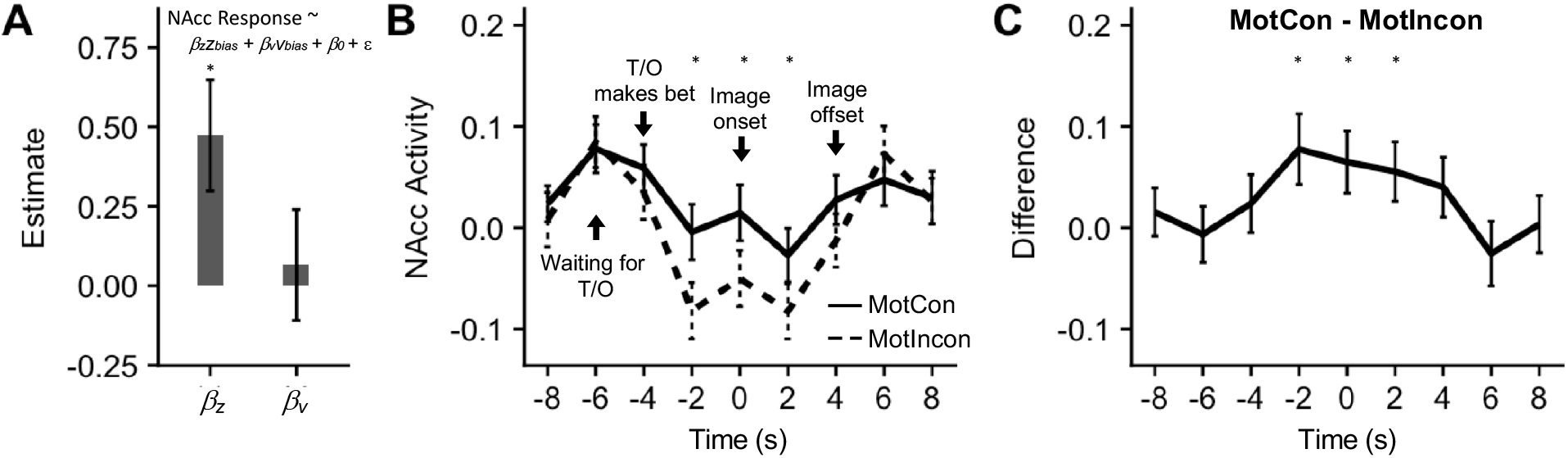
NAcc activation is associated with response bias. Linear regression predicting participants’ NAcc response from the model estimates of their starting point bias (*z_bias_*) and drift bias (*v_bias_*). The regression coefficient for *z_bias_* was significant but that for *v_bias_* was not. **B.** NAcc timecourse time-locked to image onset, corrected for hemodynamic lag by shifting the BOLD data by 4 seconds. The trial starts with the “Waiting for Teammate/Opponent” screen at −6s. The teammate or opponent makes a bet at −4s, which remains on the screen for 4s. The image is presented at 0s and stays on screen for 4s. NAcc activity was significantly higher on Motivation Consistent trials than Motivation Inconsistent trials from 2s before image onset until image offset. Solid lines: Motivation Consistent trials. Dashed lines: Motivation Inconsistent trials. **C.** Difference in activity between Motivation Consistent and Motivation Inconsistent trials peaked before image offset. *: p < 0.05, additional statistics reported in Table S2.

NAcc activity can lead to a response bias by increasing the readiness to make a particular response. This account would predict that the increase in NAcc activity was preparatory in nature and would precede the onset of the image. To test this prediction, we examined the average activity in the NAcc as a trial unfolded, separately for trials on which participants made motivation consistent categorizations (*Motivation Consistent* trials) and for trials on which participants made motivation inconsistent categorizations (*Motivation Inconsistent* trials). Consistent with a preparatory account, NAcc activity was significantly higher on Motivation Consistent trials *prior* to the image appearing on screen and remained significantly higher until image offset (Fig. 6B-C, Table S2).

These results also provide evidence against the alternative account that NAcc activation reflects the reward participants’ experience upon seeing the category they were motivated to see, as the increase in NAcc activity occurred before participants saw the image. Instead, they suggest that NAcc activity *predisposes*participants to categorize an image as the category they were motivated to see, and sustained NAcc activation increases the likelihood of making motivation consistent responses.

### Face and scene selective neural activity is associated with perceptual bias

Face and scene selective activity in the ventral occipito-temporal cortex provides a proxy measure of participants’ perception. We thus examined whether motivation affected perception by assessing if the motivation to see faces or scenes modulated this activity. We applied multivariate pattern analysis to the BOLD data to quantify the level of face and scene selective activity on each trial. Specifically, we trained a logistic regression classifier to estimate the probability that participants were seeing a scene rather than a face based on the pattern of activity in the ventral occipito-temporal cortex (see Methods).

As the proportion of scene in an image increased, the classifier predicted that the participants were seeing a scene with higher probability, indicating that the classifier tracked the amount of scene in the presented image (b = 0.121, SE = 0.005, *t(4756)* = 25.8, *p* < 0.001). There was a significant Bet x Condition interaction on classifier probability, such that the classifier was more likely to predict that participants were seeing a scene when they were motivated to see a scene than when they were motivated to see a face (b = 0.04, SE = 0.02, *t(4756)* = 2.05, *p* = 0.040; Fig. S7), indicating that the motivation to see a category increased the level of sensory evidence for that category in the visual pathway. In other words, motivation not only biased participants’ categorization of an image, it also biased their neural representation of the image.

Next, we examined how the bias in category-selective activity relates to the bias in participants’ categorical judgments (i.e. the motivational effect on a participant’s psychometric function). There was a significant triple interaction between behavioral bias, Condition and Bet on the level of category-selective activity (b = 0.057, SE = 0.017, *t(4752)* = 3.37, *p* = 0.001). To better interpret the directionality of the interaction, we performed a median split to divide participants into those with higher behavioral bias and those with lower behavioral bias. Motivation biased the classifier probability of high bias participants (b = 0.07, SE = 0.03, *t(2378)* = 2.96, *p* = 0.003), but not low bias participants (b = −0.002, SE = 0.03, *t(2373)* = −0.06, *p*= 0.953; interaction by group: *b* = 0.08, SE = 0.03, *t(4752)* = 2.14, *p* = 0.033; Fig. 7A).

**Figure 7.**
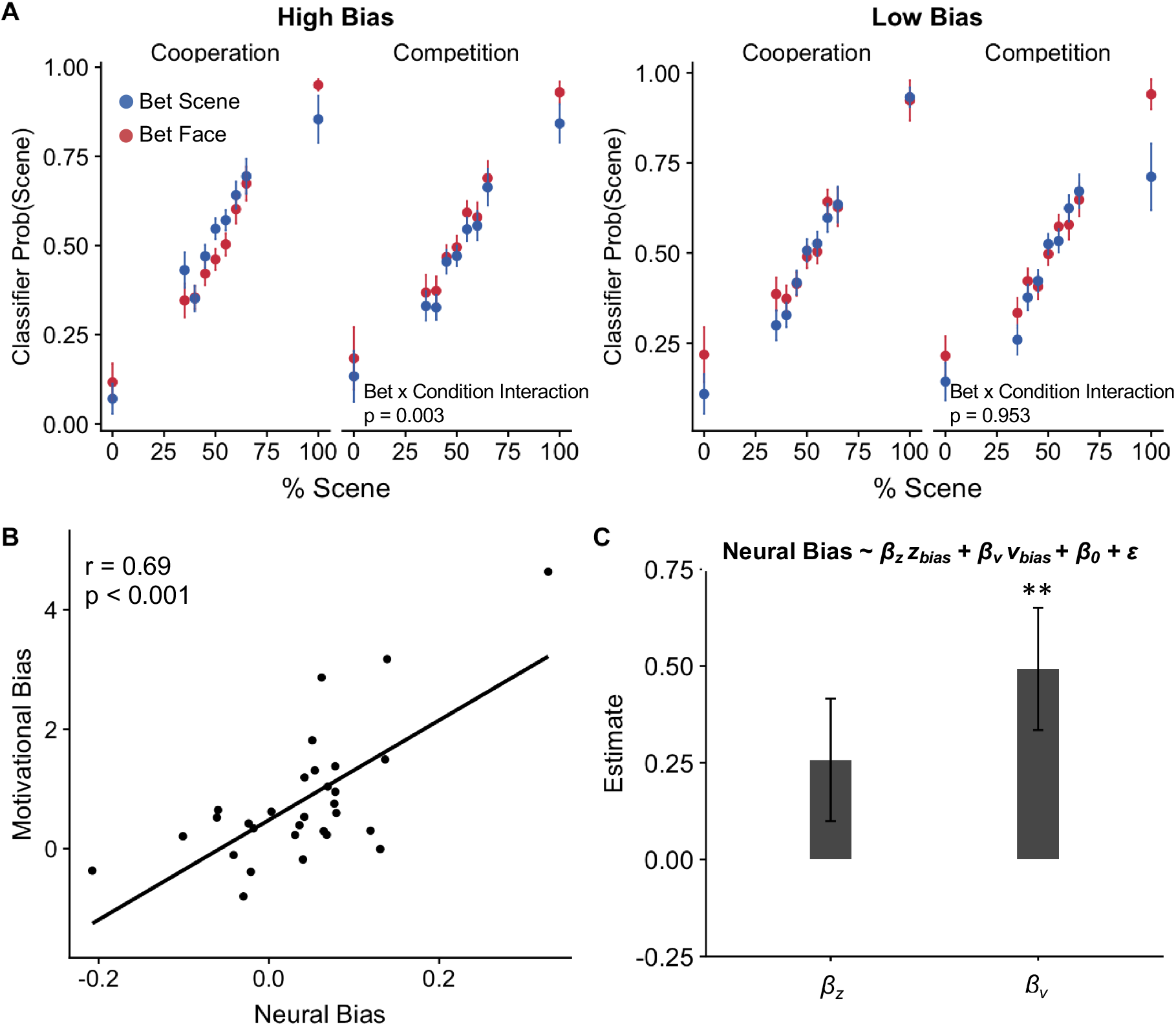
Motivation biases face and scene selective neural activity during visual categorization. **A.** Classifier probability that the presented image was a scene rather than a face based on the BOLD response in the ventral visual stream, separately for participants with high and low behavioral bias. Blue dots: teammate or opponent betting that the next image will be more scene; Red dots: teammate or opponent betting that the next image will be more face. For High Bias participants, scene probability was higher when participants were motivated to see a scene (i.e. teammate bets scene or opponent bets face) than when participants were motivated to see a face (i.e. teammate bets face or opponent bets scene). There was no effect of motivation in Low Bias participants. **B.** The effect of motivation on classifier probability (*Neural Bias*) was correlated with the extent to which a participant was biased in his or her categorizations (*Motivational Bias*). **C.** Regression coefficients of the response bias (βz) and perceptual bias (βv) when both were entered into the same model to predict participants’ Neural Bias. Only the perceptual bias was significantly associated with participants’neural bias. ** p < 0.01.

We then extracted the random slopes of the Bet x Condition interaction on classifier probability to obtain a measure of the extent to which motivation biased face and scene selective activity in each participant. The bias in participant’s face and scene selective activity correlated strongly with their behavioral bias (r = 0.69, p < 0.001, Fig. 7B), indicating that participants who were more biased in their categorizations were also more biased their neural representation of the presented image.

We then sought to relate the bias in face and scene selective activity to response and perceptual biases more specifically. When model estimates of the two biases were entered as predictors in the same linear regression model, participants’ perceptual bias was associated with the bias in face and scene selective neural activity (β_v_ = 0.49, SE = 0.16, *t*(27) = 3.12, p =0.004), but their response bias was not (β_z_ = 0.26, SE = 0.16, t(27) = 1.63, p = 0.115, Fig. 7C).

Together with our earlier analyses on NAcc activity, these results suggest distinct neural contributions to participants’ biased categorizations. While the NAcc was associated with a response bias, the modulation of face and scene selective activity in visual areas was associated with a perceptual bias. By combining computational modeling with neuroimaging, we identified two dissociable neurocomputational components underlying motivational biases in perceptual judgments.

## Discussion

This study combines computational modeling of behavior and fMRI to examine whether and why people exhibit biases towards seeing what they want to see. In a novel behavioral paradigm, we demonstrated that people indeed make biased perceptual judgments, more often labeling ambiguous images as corresponding to a reward-associated category. This is true even though participants were incentivized to accurately report their perceptual experience, and thus earned less money in the experiment when making biased judgments. Evidence from computational modeling suggests that motivational effects on perceptual judgments could be attributed to both a response bias and a bias in perceptual processing. While the response bias was associated with anticipatory activity in the nucleus accumbens (NAcc), the bias in perceptual processing was associated with the modulation of category-selective neural activity in the ventral visual stream. These results provide converging evidence for two distinct contributions to motivational influences on perceptual judgments, and shed light on the neurocomputational mechanisms underlying each bias.

The claim that perceptual processes are influenced by motivational factors can be traced back to the “New Look” movement in psychology, which argued that the perception of external stimuli is subject to the constant influence of a perceiver’s internal goals and states^28,29^. Recent evidence supporting this view includes studies demonstrating that perceptually ambiguous stimuli are more likely to be seen as the percept associated with favorable outcomes^4,30^, desirable objects are judged nearer than undesirable ones^31^, and desirable food items are judged as larger by dieters than non-dieters^32^. Whether these results reflect a bias in subjective reports or a bias in perception remains a topic of intense debate (see open peer commentary for ^7^). In particular, since these studies rely primarily on subjective reports, and participants often have an incentive to report seeing what they want to see, there is reason to suspect that subjective reports might not reflect one’s underlying perceptual experience.

Our work builds on earlier efforts to understand motivated perception using computational models. An earlier study^33^ similarly decomposed motivational biases into response and perceptual components by mapping them respectively onto biases in the starting point and drift rate of a drift diffusion model. There are, however, non-perceptual explanations for a bias in drift rate. For example, a response bias that becomes stronger over the course of a trial (i.e. a *dynamic* response bias) could also result in a biased drift rate^34,35^. As such, a bias in drift rate alone is insufficient evidence of a bias in perception.

A strength of the current work is that in addition to modeling the behavioral data using a drift diffusion model, we used functional neuroimaging to measure the level of sensory evidence in visual areas of the brain. We found that motivation enhances the sensory evidence of the motivationally consistent category in the ventral visual stream, and the degree of sensory enhancement was associated with the bias in drift rate but not the bias in starting point. In relating the neural data to the modeling results, we achieve two complementary goals – (i) the neural results provide convergent validity of our interpretation of the bias in drift rate as a perceptual bias, (ii) the modeling results provide a computational description of how the bias in sensory activity contributes to biases in perceptual judgments.

Perceptual judgments are thought to be computed by comparing the activity of neurons selective to different perceptual features^19,20^. Within this framework, the nervous system “reads out” the activity of face-selective and scene-selective neurons as sensory evidence for faces and scenes respectively. A perceptual judgment can then be determined by comparing the activity of face-selective and scene-selective neurons. Our results indicate that motivation biases this comparison by enhancing the activity of the neurons selective to the category participants were motivated to see. This enhancement could in turn reflect the biased processing of incoming sensory information, with the biasing signal originating from frontoparietal attention regions^36^.

Indeed, we found that the intraparietal sulcus (IPS) and frontal eye fields (FEF) were more active when participants made motivationally consistent judgments. The IPS and FEF are part of the dorsal attention network associated with the top-down control of attention^37,38^. Their involvement in our task suggest that the bias in perceptual processing might be in part mediated by dynamic changes in the focus of attention^39^. In addition to the frontoparietal activations, the dorsal anterior cingulate cortex (dACC) and insula were also more active on Motivation Consistent trials. The dACC and insula are part of a salience network involved in the detection of motivationally salient stimuli^40,41^, and the dACC has been recently implicated in determining what stimulus feature to attend to in a perceptual decision-making task^42^. The increased activity in the salience network on Motivation Consistent trials might be responsible for the selection of motivationally relevant features for enhanced processing. However, this interpretation is speculative, and future studies will be needed to clarify the role of each region in biasing perceptual judgments.

On the other hand, participants’ response bias was associated with activity in the nucleus accumbens (NAcc). This is consistent with behavioral neuroscience work suggesting that dopaminergic projections to the NAcc biases animals towards responses associated with greater reward^12–14,16^. Both human neuroimaging and animal physiology studies have also shown that the NAcc is activated in anticipation of reward^10,11^. Our results suggest a functional role for this anticipatory activity. In particular, they suggest that the NAcc increases participants’ readiness to respond in a motivation consistent manner. When the motivation consistent response is aligned with task demands (e.g., pressing a lever for reward), this preparatory response facilitates faster responding for reward^15,43^. However, when the motivation consistent response conflicts with task demands, as was the case in our task, the preparatory response is maladaptive and impairs performance on the task (see also ^44^).

Our results add to the rich literature on perceptual decision-making in cognitive neuroscience by dissociating motivation (“wanting to see”) from optimal task performance (“reward-maximization”). Previous studies have manipulated the reward associated with different perceptual alternatives, and found that reward biases responses but not perceptual processing^45–48^. We speculate that these results differ markedly from ours because our paradigm tapped into distinct biasing mechanisms. In these earlier studies, participants would earn the additional reward if they correctly categorized the stimulus as the rewarded category. Under this payoff scheme, biasing responses towards the option associated with larger reward would result in greater cumulative reward over the course of the experiment^45,49^. Hence, the bias in these earlier experiments likely reflects a strategic shift in responses to maximize reward on the task, and thus not affect sensory processing.

By contrast, in our task, the additional reward associated with the motivationally consistent category was independent of participants’responses. For example, if the teammate bet that the next image would have more face, participants would receive the bonus if the upcoming image indeed had more face, regardless of how they responded on the trial. In this case, a bias towards the motivationally consistent category would lower participants’ earnings by hurting their accuracy on the categorization task. Thus, the biases observed in our task cannot be explained by existing normative models of judgment and decision-making that assume organisms adjust their choice strategies to maximize expected reward. Instead, they highlight a motivational component to perceptual judgments – wanting an outcome to be true can impinge on one’s perceptual judgment, even when doing so could lead to lower rewards in the long run. Our results suggest that this bias reflects not only a response bias, but also a perceptual bias.

At a broader level, this work provides a novel bridge between social psychology and cognitive neuroscience. Using tools and analytical techniques from cognitive neuroscience, we examine the neurocomputational mechanisms underlying an age-old phenomenon of interest in social psychology. In doing so, we offer a fresh perspective on a classic debate. Unlike earlier work that assesses *whether* motivation biases perception, we provide a neurocomputational account of *how* motivation biases perception. The results also complement the existing literature on motivated person perception, which has focused primarily on the neural and computational mechanisms by which people form overly positive evaluations of themselves and close others^50–52^. Our work extends the phenomenon beyond the domain of social attributions, and show that motivated visual perception can be similarly characterized as a change in initial beliefs (i.e. starting point) and information updating (i.e. drift rate).

Desires and wants exert a powerful influence over how people make sense of the world. Recent studies have examined the neural mechanisms underlying motivational biases across a variety of human reasoning and evaluative processes^53^, including how the brain learns more from positive outcomes than negative ones^54^ and why people form unrealistically optimistic expectations about future events^55^. Here, we demonstrate that motivation biases human cognition as early as visual perception, and provide a neurocomputational account of this effect. The current work extends our understanding of motivational biases and provides a starting point to explore how motivation acts on different neural systems at different stages of information processing to influence human cognition.

## Methods

### Participants

Thirty-three participants were recruited from the Stanford community, and provided written, informed consent prior to the start of the study. All experimental procedures were approved by the Stanford Institutional Review Board. Participants were paid between $30-$50 depending on their performance on the task. Data from three participants were discarded because of excessive head motion (> 3mm) during one or more scanning sessions, yielding an effective sample size of thirty participants (17 male, 13 female, ages 18-43, mean age = 22.3).

### Stimuli

For each participant, seven sets (one for the practice task and six for the experimental task) of composite stimuli were created. Each stimulus set consists of 40 grey-scale images, each comprising a mixture of a face image and a scene image in varying proportions (1 x 100% scene, 3 x 65% scene, 5 x 60% scene, 7 x 55% scene, 8 x 50% scene, 7 x 45% scene, 5 x 40% scene, 3 x 35% scene, 1 x 0% scene). Scene images comprised of half indoor scenes and half outdoor scenes, while face images comprised of half male faces and half female faces. All faces were frontal photographs posing a neutral expression, and were taken from the Chicago Face Database^56^. Stimuli were presented using MATLAB software (MathWorks) and the Psychophysics Toolbox^57^.

### Practice Task

Participants first performed 40 practice trials in which they were presented with composite Face/Scene images (see *Stimuli*). Each image was presented for four seconds, during which participants had to judge whether the image contained a greater proportion of face (“more face”) or a greater proportion of scene (“more scene”). Participants earned 10 cents for each correct categorization. They then indicated how confident they were in their classification on a 1 to 5 scale. If they did not respond within four seconds, the trial timed out and they would not earn a bonus on that trial. After a variable inter-trial interval (ITI, 2s-4s), they moved on to the next trial. We collected participants’ anatomical scans while they performed the practice task.

### Experimental Task

The experimental task consists of four fMRI runs, each approximately 8 minutes long. Participants performed two runs of the Cooperation condition and two runs of the Competition condition (interleaved order, counterbalanced across participants, Fig. 1A). Each run consisted of 40 trials. In the Cooperation condition, participants were told that they would perform a visual categorization task with a teammate. At the start of each trial, their teammate would make a bet on the image type of the upcoming image (“more face” or “more scene”, presented for 4 seconds). Participants were then presented with a composite image created by averaging a face image and a scene image in different proportions (see *Stimuli*). If the teammate’s bet was correct, both the teammate and participants would earn 40 cents. If the teammate’s bet was wrong, both the teammate and participants would lose 40 cents.

Participants then had four seconds to make a categorization on whether the image contained “more face” or “more scene” (see also *Practice Task*). Participants earned 10 cents for each correct categorization. They then indicated how confident they were in their classification on a 1 to 5 scale. If they did not respond within four seconds, the trial timed out and they would not earn a bonus on that trial (though the bet would still be implemented). After a variable ITI (2s-4s), they moved on to the next trial. In the Competition condition, participants performed the task with an opponent. The trial structure was identical to the Cooperation condition, except that if the opponent’s bet was correct, the opponent would earn 40 cents while participants lose 40 cents. If the opponent’s bet was wrong, the opponent loses 40 cents while participants earn 40 cents. As such, participants were motivated to see the image type their teammate bet on, and to see the image type opposite of what their opponent bet on. Participants were told that the teammate and opponent would not be informed about their responses, and would thus not be motivated by a desire to appease or defy their teammate or opponent.

Crucially, the outcome of the bets was contingent on whether the image objectively contained more face or more scene, and was not contingent on participants’ subjective categorization. Hence, the reward maximizing strategy was to ignore the bets and categorize the images as accurately as possible (Fig. 1B). Bets by both the teammate and the opponent were pseudo-randomized such that they were accurate on exactly 50% of the trials. As such, participants’ earnings in the experiment depended solely on their performance on the categorization task. We computed participants’ performance as the average number of correct categorizations.

### Localizer Task

To identify BOLD activation associated with viewing faces or scenes, we had participants perform a localizer task at the end of the experiment. Participants viewed 5 blocks of 15 unambiguous Faces and 5 blocks of 15 unambiguous Scenes (blocks were interleaved, and order was counterbalanced across participants). In Face blocks, participants were sequentially presented with face images, and had to indicate whether each face was male or female. In Scene blocks, participants were sequentially presented with scene images, and had to indicate whether each scene was indoors or outdoors. Each image was presented for 2 seconds, with a 2-second ITI. Participants took a self-timed break between blocks. The localizer task was split into 2 scans.

### fMRI data acquisition and preprocessing

MRI data were collected using a 3T General Electric MRI scanner. Functional images were acquired in interleaved order using a T2*-weighted echo planar imaging (EPI) pulse sequence (46 transverse slices, TR=2s, TE=25ms, flip angle=77°, voxel size 2.9 mm^3^). Anatomical images were acquired at the start of the session with a T1-weighted pulse sequence (TR = 7.2ms, TE = 2.8ms, flip angle=12°, voxel size 1 mm^3^). Image volumes were preprocessed using FSL/FEAT v.5.98 (FMRIB software library, FMRIB, Oxford, UK). Preprocessing included motion correction, slice-timing correction, removal of low-frequency drifts using a temporal high-pass filter (100ms cutoff), and spatial smoothing (4-mm FWHM). For multivoxel classification analyses, we trained and tested our classifier in each participant’s native space. For all other analyses, functional volumes were first registered to participants’ anatomical image (Boundary-Based Registration) and then to a template brain in Montreal Neurological Institute (MNI) space (affine transformation with 12 degrees of freedom).

### Psychometric functions

We modeled participants’ behavioral data using generalized linear mixed effects models (GLMM), which allows for the modeling of all of the data in one step rather than fitting a separate model for each participant^58^. Three separate linear mixed effects model were fit to participants’ data to estimate the effect of the motivation manipulation (i.e. the “Bet” by the teammate or opponent) on participants’ categorizations. The first two models were fit to data from the Cooperation and Competition condition to estimate the effect of the teammate and opponent’s bet respectively, while the third model was fit to data from both conditions to estimate the Bet x Condition interaction. All models included random intercepts and random slopes for the effects of the teammate or opponent’s bet (scene/face) to account for random variability across participants. The third model also included random intercepts and slopes for the Condition and the Condition x Bet interaction. Models were estimated using the glmer function in the lme4 package in R^59^, with p-values computed from t-tests with Satterthwaite approximation for the degrees of freedom as implemented in the lmerTest package^60^. The estimates of the random slope of the interaction term reflected the extent to which each participant’s categorizations were biased by the motivation manipulation.

### Reaction time analyses

We ran a series of linear mixed effects models to examine the effect of motivation on the pattern of reaction times. We first examined whether reaction times were faster when making motivation consistent responses (e.g., responding face when motivated to see more face) than when making motivation inconsistent responses (e.g., responding face when motivated to see more scene), controlling for the absolute difference between percent scene and percent face as a measure of stimulus uncertainty. Next, we examined motivational effects on reaction times separately for trials on which participants responded face and those on which they responded scene. For each group, we tested if the reaction times differed depending on the category participants were motivated to see, again controlling for stimulus uncertainty. To visualize these results, we plot the average reaction time for face and scene responses at each level of % scene, separately for when participants were motivated to see more face and when participants were motivated to see more scene. Finally, we repeated these analyses separately for the Cooperation and Competition condition, and also examined whether motivational effects on reaction time differed between the two conditions by testing for a triple interaction of Condition x Motivation x Response on reaction times.

### Robust Regression Analysis

To examine the relationship between motivational bias and task performance, we fit a linear model by robust regression. Model fitting was performed using the rlm function from the “MASS” package in R. Robust regression is an alternative to linear regression that is less sensitive to outliers^61^. Statistical significance of the regression coefficient was assessed by performing a robust F test.

### Drift Diffusion Model

The drift diffusion model assumes that decisions are made by accumulating evidence over time until it crosses one of two decision bounds^9^ (Fig. 3A). The starting point and rate of evidence accumulation were determined by free parameters *z* and *v* respectively. The distance between the two boundaries depended on free parameter *a*, while time not related to decision process (e.g., stimulus encoding, motor response) was modeled by the free parameter *t*.

Model parameters were estimated from participants’ categorizations and RT distributions using hierarchical Bayesian estimation as implemented by the HDDM toolbox^25^. Parameters for individual participants were assumed to be randomly drawn from a group-level distribution. In the fitting procedure, each participant’s parameters both contributed to and were constrained by the estimates of group-level parameters. Markov chain Monte Carlo (MCMC) sampling methods were used to estimate the joint posterior distribution of all model parameters (100,000 samples; burn-in = 10,000 samples; thinning = 2). We estimated both group-level parameters as well as parameters for each individual participant, which allowed us to examine biases in both the entire sample and in each individual participant. To account for outliers generated by a process other than that assumed by the model (e.g., lapses in attention, accidental button press), we estimated a mixture model where 5% of trials were assumed to be distributed according to a uniform distribution.

In the *z & v* model, we modeled *z* as a function of the motivation consistent category and an intercept term. The HDDM package implements *z* as the *relative* starting point, bound between 0 and 1, with 0.5 reflecting an unbiased starting point. As such, we used the inverse logit link function to restrict *z* to values between 0 and 1:

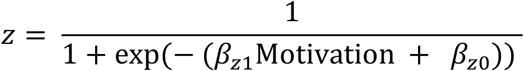

where *Motivation* denotes the motivation consistent category, *β_z1_* reflects the effect of motivation on the starting point (*z_bias_*) and *β_z0_* denotes an intercept term. We modeled *v* as a linear combination of the category participants were motivated to see, the level of percentage scene and an intercept term:

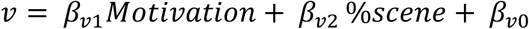

where *β_v1_* reflects the effect of motivation on the drift rate (*v_bias_*) and *β_v2_* reflects the effect of percentage scene on the drift rate. We demeaned % scene prior to entering it into the model such that the intercept term, *β_v0_*, would also reflect the intrinsic drift bias. For each of the bias parameters (*z_bias_* and *v_bias_*), we computed the proportion of posterior samples that were greater than 0 (Fig. 3C and D). We consider the estimates to be credibly above zero if more than 95% of the posterior distribution were to be greater than zero.

To examine if either of the biases were sufficient for explaining the data, we fit two additional comparison models in which only *z*(*z* model) or only *v*(*v* model) was biased by motivation. In the *z* model, *β_z1_* was fixed at 0 while in the *v* model, *β_v1_* was fixed to zero. As a baseline for comparison, we also fit a *null* model in which neither the starting point nor drift rate were biased by motivation. We then compared the four models using deviance information criterion (DIC), which is a measure of model performance that appropriately penalizes for model complexity in hierarchical models^26^. To verify that DIC is an accurate metric for model comparison, we also ran a model recovery study to examine if DIC correctly recovers the model used to generate simulated data (see Supplemental Note)

Model convergence of all models was formally assessed using the Gelman-Rubin 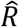 statistic^62^, which runs multiple Markov chains to compare within-chain and between-chain variances. Large differences 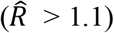 between these variances would signal non-convergence. In addition, we examined each trace to check that there were no drifts or large jumps, which would also suggest non-convergence. We report model convergence metrics, posterior means and 95% credible intervals of all parameters in Table S1.

### Model simulations

We generated 500 simulated datasets, each comprising the same number of participants performing the same number of trials as the real dataset. Each dataset was generated with parameter values sampled from the posterior distribution estimated by the *z & v* model. These datasets reflect the pattern of choice and reaction time data if participants’ behavior were perfectly described by the model. To compare the simulations to real data, we averaged over the 500 simulations to obtain the average reaction time of face and scene responses at each level of % scene, separately for when participants were motivated to see more face and when participants were motivated to see more scene.

Given that the drift diffusion model was fitted to reaction time distributions (rather than the mean), we also assessed how well the model accounts for the shape of participants’ reaction time distributions. For each participant, we overlay the distribution of simulated reaction times with the true reaction time distributions, separately for face and scene responses. These plots serve as posterior predictive checks to assess how well the model aligns with participants data.

### GLM

We implemented a linear model (GLM 1) to contrast BOLD activity on Motivation Consistent trials and that on Motivation Inconsistent trials. A Motivation Consistent trial was defined as a trial on which participants categorized an image as the category they were motivated to see. This contrast would thus identify voxels in the brain which were significantly more active when participants reported seeing what they wanted to see, versus what they did not want to see. Stimulus onset, the objective % scene, reaction time and head movement parameters were included as nuisance regressors. With the exception of head movement parameters, all regressors were convolved with a hemodynamic response function. The GLM was estimated throughout the whole brain using FSL/FEAT v.5.98 available as part of the FMRIB software library (FMRIB). Correction for multiple comparisons was performed using threshold free cluster enhancement (TFCE) with an alpha of 0.05, as implemented by the *randomise* tool in FSL^27^.

We implemented a second linear model (GLM 2) in which the onset of each trial was modeled as a separate regressor. This model allowed us estimate a separate statistical map for each trial (i.e. single trial activation patterns). We then used these maps as inputs to the multivoxel pattern analyses (see below). As was the case in GLM 1, reaction time and head movement parameters were included as nuisance regressors.

### NAcc region of interest analyses

We defined an independent NAcc region of interest (ROI) using the Harvard-Oxford subcortical structural atlas (available for download at https://neurovault.org/collections/EAAXGDRJ/). For each participant, we extracted the average z-statistic of the Motivation Consistent > Motivation Inconsistent contrast (GLM 1) within the NAcc ROI. This average z-statistic reflects the extent to which an ROI is more reliably active on Motivation Consistent trials than on Motivation Inconsistent trials for that participant, and was taken as the participant’s NAcc response. The NAcc response was then regressed against estimates of starting point and drift bias estimated by the *z & v* model (see *Relating model parameters to behavior and neural measures*).

To examine how the NAcc response unfolded across a trial, we extracted and z-scored the mean timecourse in the NAcc ROI from each run. Each timecourse was shifted by 2 TRs (4 seconds) to correct for hemodynamic lag. We extracted the data from 8 seconds before stimulus onset to 8 seconds after stimulus onset to obtain the timecourse of a single trial, and computed the average timecourse of activity separately for Motivation Consistent trials and Motivation Inconsistent trials. At each time point, we assessed if activity was higher on Motivation Consistent trials and than on Motivation Inconsistent trials using a paired sample t-test.

### Multivoxel pattern analyses

Multivoxel pattern analyses were performed using tools available as part of the *nilearn* Python module^63^. An L1-regularized logistic regression model (C = 1) was trained on BOLD data from the localizer task to classify the image category participants were seeing on each localizer trial. Notably, the localizer task was unrelated to Face vs. House discrimination (see Localizer Task). Hence, the category-selective patterns identified by the classifier were specific to seeing faces or scenes, and not confounded by response-related information.

Analysis was restricted to voxels in a ventral visual stream mask consisting of the bilateral occipital lobe and ventral temporal cortex. The ventral occipito-temporal regions of the brain are thought to be important in perceiving object categories such as faces and scenes^22^. The mask was created in MNI space using anatomical masks defined by the Harvard-Oxford Cortical Structural Atlas. The mask was then transformed into each participant’s native space using FSL’s FLIRT implementation, and classification was performed in participants’ native space.

The trained model was then applied to the single trial activation patterns in the experimental task (GLM 2). On each trial, the classifier returned the probability that the participant was seeing a scene rather than a face based on activity in the ventral visual stream mask. We then modeled classifier probability on each trial using a linear mixed effects model with the percentage scene of an image, the task Condition (Cooperation/Competition), the teammate or opponent’s Bet (Face/Scene) and the interaction between Condition and Bet as predictor variables. The models included random intercepts and random slopes for each of the predictor variables to account for the random variability across participants. The estimate of the random slope of the interaction term of a participant reflected the extent to which classifier probability was biased by the motivation manipulation for that particular participant. This estimate was taken as a measure of neural bias, that is, the extent to which category-selective activity in the ventral visual stream for that particular participant was modulated by motivation.

### Relating model parameters to behavior and neural measures

We used linear regression to examine the relationship between model parameters and neural activity. We entered participant-level estimates of the starting point bias (*z* bias) and the drift bias (*v* bias) as predictor variables in regression models. The first model was used to predict participants’ NAcc response to the Motivation Consistent – Motivation Inconsistent contrast (GLM 1), and assessed the extent to which each bias was associated with NAcc activity. The second model was used to predict participants’ neural bias, and assessed the extent to which each bias contributed to the modulation of category-selective activity in the ventral visual stream.

## Data and code availability

Behavioral data of both the reported experiment as well as the inlab replication are available at: https://github.com/ycleong/MotivatedPerception. Custom code for modeling and neuroimaging analyses are available at the same repository. Unthresholded p-map of the Motivation Consistent – Motivation Inconsistent contrast is available at: https://neurovault.org/collections/EAAXGDRJ/images/62743/. Raw neuroimaging data available on request.

## Acknowledgments

We thank Ian Ballard and members of the Stanford Social Neuroscience Laboratory for scientific discussions and helpful comments on earlier versions of the manuscript. The research was supported by the Stanford Neuroscience Institute NeuroChoice Initiative.

## Author contributions

Y.C.L., B.L.H., and J.Z. designed the study; Y.C.L. and Y.W. collected and analyzed the data; Y.C.L. and J.Z. wrote the manuscript, with revisions from Y.W. and B.L.H.

## References

1. Bruner, J. S. & Goodman, C. C. Value and need as organizing factors in perception. J. Abnorm. Soc. Psychol. 42, 33–44 (1947).

2. Dunning, D. & Balcetis, E. Wishful Seeing: How Preferences Shape Visual Perception. Curr. Dir. Psychol. Sci. 22, 33–37 (2013).

3. Hastorf, A. H. & Cantril, H. They saw a game; a case study. J. Abnorm. Soc. Psychol. 49, 129–134 (1954).

4. Balcetis, E. & Dunning, D. See what you want to see: Motivational influences on visual perception. J. Pers. Soc. Psychol. 91, 612–625 (2006).

5. Kunda, Z. The case for motivated reasoning. Psychol. Bull. 108, 480–498 (1990).

6. Goldiamond, I. Indicators of perception: I. Subliminal perception, subception, unconscious perception: An analysis in terms of psychophysical indicator methodology. Psychol. Bull. 55, 373–411 (1958).

7. Firestone, C. & Scholl, B. J. Cognition does not affect perception: Evaluating the evidence for ‘top-down’ effects. Behav. Brain Sci. N. Y. 39, (2016).

8. Forstmann, B. U., Ratcliff, R. & Wagenmakers, E.-J. Sequential Sampling Models in Cognitive Neuroscience: Advantages, Applications, and Extensions. Annu. Rev. Psychol. 67, 641–666 (2016).

9. Ratcliff, R. & McKoon, G. The Diffusion Decision Model: Theory and Data for Two-Choice Decision Tasks. Neural Comput. 20, 873–922 (2008).

10. Berridge, K. C. The debate over dopamine’s role in reward: the case for incentive salience. Psychopharmacology (Berl.) 191, 391–431 (2007).

11. Knutson, B., Adams, C. M., Fong, G. W. & Hommer, D. Anticipation of Increasing Monetary Reward Selectively Recruits Nucleus Accumbens. J. Neurosci. 21, RC159–RC159 (2001).

12. Floresco, S. B. The Nucleus Accumbens: An Interface Between Cognition, Emotion, and Action. Annu. Rev. Psychol. 66, 25–52 (2015).

13. Ikemoto, S. & Panksepp, J. The role of nucleus accumbens dopamine in motivated behavior: a unifying interpretation with special reference to reward-seeking. Brain Res. Rev. 31, 6–41 (1999).

14. Nicola, S. M. The nucleus accumbens as part of a basal ganglia action selection circuit. Psychopharmacology (Berl.) 191, 521–550 (2007).

15. McGinty, V. B., Lardeux, S., Taha, S. A., Kim, J. J. & Nicola, S. M. Invigoration of Reward Seeking by Cue and Proximity Encoding in the Nucleus Accumbens. Neuron 78, 910–922 (2013).

16. Stopper, C. M. & Floresco, S. B. Contributions of the nucleus accumbens and its subregions to different aspects of risk-based decision making. Cogn. Affect. Behav. Neurosci. 11, 97–112 (2011).

17. Gold, J. I. & Shadlen, M. N. The Neural Basis of Decision Making. Annu. Rev. Neurosci. 30, 535–574 (2007).

18. Heekeren, H. R., Marrett, S. & Ungerleider, L. G. The neural systems that mediate human perceptual decision making. Nat. Rev. Neurosci. 9, 467–479 (2008).

19. Shadlen, M. N., Britten, K. H., Newsome, W. T. & Movshon, J. A. A computational analysis of the relationship between neuronal and behavioral responses to visual motion. J. Neurosci. 16, 1486–1510 (1996).

20. Heekeren, H. R., Marrett, S., Bandettini, P. A. & Ungerleider, L. G. A general mechanism for perceptual decision-making in the human brain. Nature 431, 859 (2004).

21. Summerfield, C. & Egner, T. Expectation (and attention) in visual cognition. Trends Cogn. Sci. 13, 403–409 (2009).

22. Grill-Spector, K. The neural basis of object perception. Curr. Opin. Neurobiol. 13, 159–166 (2003).

23. Hasson, U., Hendler, T., Bashat, D. B. & Malach, R. Vase or Face? A Neural Correlate of Shape-Selective Grouping Processes in the Human Brain. J. Cogn. Neurosci. 13, 744–753 (2001).

24. White, C. N. & Poldrack, R. A. Decomposing bias in different types of simple decisions. J. Exp. Psychol. Learn. Mem. Cogn. 40, 385–398 (2014).

25. Wiecki, T. V., Sofer, I. & Frank, M. J. HDDM: Hierarchical Bayesian estimation of the Drift-Diffusion Model in Python. Front. Neuroinformatics 7, (2013).

26. Spiegelhalter, D. J., Best, N. G., Carlin, B. P. & Van Der Linde, A. Bayesian measures of model complexity and fit. J. R. Stat. Soc. Ser. B Stat. Methodol. 64, 583–639 (2002).

27. Smith, S. M. & Nichols, T. E. Threshold-free cluster enhancement: Addressing problems of smoothing, threshold dependence and localisation in cluster inference. NeuroImage 44, 83–98 (2009).

28. Allport, F. H. Theories of perception and the concept of structure: A review and critical analysis with an introduction to a dynamic-structural theory of behavior. (John Wiley & Sons Inc, 1955). doi:10.1037/11116-000

29. Bruner, J. S. On perceptual readiness. Psychol. Rev. 64, 123–152 (1957).

30. Balcetis, E., Dunning, D. & Granot, Y. Subjective value determines initial dominance in binocular rivalry. J. Exp. Soc. Psychol. 48, 122–129 (2012).

31. Balcetis, E. & Dunning, D. Wishful Seeing: More Desired Objects Are Seen as Closer. Psychol. Sci. 21, 147–152 (2010).

32. van Koningsbruggen, G. M., Stroebe, W. & Aarts, H. Through the eyes of dieters: Biased size perception of food following tempting food primes. J. Exp. Soc. Psychol. 47, 293–299 (2011).

33. Voss, A., Rothermund, K. & Brandtstädter, J. Interpreting ambiguous stimuli: Separating perceptual and judgmental biases. J. Exp. Soc. Psychol. 44, 1048–1056 (2008).

34. Moran, R. Optimal decision making in heterogeneous and biased environments. Psychon. Bull. Rev. 22, 38–53 (2015).

35. Hanks, T. D., Mazurek, M. E., Kiani, R., Hopp, E. & Shadlen, M. N. Elapsed Decision Time Affects the Weighting of Prior Probability in a Perceptual Decision Task. J. Neurosci. 31, 6339–6352 (2011).

36. Serences, J. T. Value-Based Modulations in Human Visual Cortex. Neuron 60, 1169–1181 (2008).

37. Corbetta, M. & Shulman, G. L. Control of goal-directed and stimulus-driven attention in the brain. Nat. Rev. Neurosci. 3, 201 (2002).

38. Petersen, S. E. & Posner, M. I. The Attention System of the Human Brain: 20 Years After. Annu. Rev. Neurosci. 35, 73–89 (2012).

39. Leong, Y. C., Radulescu, A., Daniel, R., DeWoskin, V. & Niv, Y. Dynamic Interaction between Reinforcement Learning and Attention in Multidimensional Environments. Neuron 93, 451–463 (2017).

40. Menon, V. Salience Network. in Brain Mapping (ed. Toga, A. W.) 597–611 (Academic Press, 2015). doi:10.1016/B978-0-12-397025-1.00052-X

41. Sridharan, D., Levitin, D. J. & Menon, V. A critical role for the right fronto-insular cortex in switching between central-executive and default-mode networks. Proc. Natl. Acad. Sci. 105, 12569–12574 (2008).

42. Shenhav, A., Straccia, M. A., Musslick, S., Cohen, J. D. & Botvinick, M. M. Dissociable neural mechanisms track evidence accumulation for selection of attention versus action. Nat. Commun. 9, 2485 (2018).

43. Niv, Y., Daw, N. D., Joel, D. & Dayan, P. Tonic dopamine: opportunity costs and the control of response vigor. Psychopharmacology (Berl.) 191, 507–520 (2007).

44. Krebs, R. M., Boehler, C. N., Egner, T. & Woldorff, M. G. The Neural Underpinnings of How Reward Associations Can Both Guide and Misguide Attention. J. Neurosci. 31, 9752–9759 (2011).

45. Feng, S., Holmes, P., Rorie, A. & Newsome, W. T. Can Monkeys Choose Optimally When Faced with Noisy Stimuli and Unequal Rewards? PLOS Comput. Biol. 5, e1000284 (2009).

46. Mulder, M. J., Wagenmakers, E.-J., Ratcliff, R., Boekel, W. & Forstmann, B. U. Bias in the Brain: A Diffusion Model Analysis of Prior Probability and Potential Payoff. J. Neurosci. 32, 2335–2343 (2012).

47. Rorie, A. E., Gao, J., McClelland, J. L. & Newsome, W. T. Integration of Sensory and Reward Information during Perceptual Decision-Making in Lateral Intraparietal Cortex (LIP) of the Macaque Monkey. PLOS ONE 5, e9308 (2010).

48. Summerfield, C. & Koechlin, E. Economic Value Biases Uncertain Perceptual Choices in the Parietal and Prefrontal Cortices. Front. Hum. Neurosci. 4, (2010).

49. Bogacz, R., Brown, E., Moehlis, J., Holmes, P. & Cohen, J. D. The physics of optimal decision making: A formal analysis of models of performance in two-alternative forced-choice tasks. Psychol. Rev. 113, 700–765 (2006).

50. Flagan, T., Mumford, J. A. & Beer, J. S. How Do You See Me? The Neural Basis of Motivated Meta-perception. J. Cogn. Neurosci. 29, 1908–1917 (2017).

51. Hughes, B. L. & Beer, J. S. Orbitofrontal Cortex and Anterior Cingulate Cortex Are Modulated by Motivated Social Cognition. Cereb. Cortex 22, 1372–1381 (2012).

52. Korn, C. W., Prehn, K., Park, S. Q., Walter, H. & Heekeren, H. R. Positively Biased Processing of Self-Relevant Social Feedback. J. Neurosci. 32, 16832–16844 (2012).

53. Hughes, B. L. & Zaki, J. The neuroscience of motivated cognition. Trends Cogn. Sci. 19, 62–64 (2015).

54. Lefebvre, G., Lebreton, M., Meyniel, F., Bourgeois-Gironde, S. & Palminteri, S. Behavioural and neural characterization of optimistic reinforcement learning. Nat. Hum. Behav. 1, 0067 (2017).

55. Sharot, T., Korn, C. W. & Dolan, R. J. How unrealistic optimism is maintained in the face of reality. Nat. Neurosci. 14, 1475–1479 (2011).

56. Ma, D. S., Correll, J. & Wittenbrink, B. The Chicago face database: A free stimulus set of faces and norming data. Behav. Res. Methods 47, 1122–1135 (2015).

57. Brainard, D. H. The Psychophysics Toolbox. Spat. Vis. 10, 433–436 (1997).

58. Knoblauch, K. & Maloney, L. T. Modeling Psychophysical Data in R. (Springer Science & Business Media, 2012).

59. Bates, D., Mächler, M., Bolker, B. & Walker, S. Fitting Linear Mixed-Effects Models Using lme4. J. Stat. Softw. Artic. 67, 1–48 (2015).

60. Kuznetsova, A., Brockhoff, P. B. & Christensen, R. H. B. lmerTest: Tests in Linear Mixed Effects Models. (2016).

61. Venables, W. N. & Ripley, B. D. Modern Applied Statistics with S. (Springer Science & Business Media, 2003).

62. Gelman, A. & Rubin, D. B. Inference from Iterative Simulation Using Multiple Sequences. Stat. Sci. 7, 457–472 (1992).

63. Abraham, A. et al. Machine learning for neuroimaging with scikit-learn. Front. Neuroinformatics 8, (2014).

